# Genetic diversity and structure in wild Robusta coffee (*Coffea canephora* A. Froehner) populations in Yangambi (DR Congo) and their relation with forest disturbance

**DOI:** 10.1101/2022.10.07.511267

**Authors:** Jonas Depecker, Lauren Verleysen, Justin A Asimonyio, Yves Hatangi, Jean-Léon Kambale, Ithe Mwanga Mwanga, Ebele Tshimi, Benoit Dhed’a, Yves Bawin, Ariane Staelens, Piet Stoffelen, Tom Ruttink, Filip Vandelook, Olivier Honnay

## Abstract

Degradation and regeneration of tropical forests can strongly affect gene flow in understorey species, resulting in genetic erosion and changes in genetic structure. Yet, these processes remain poorly studied in tropical Africa. *Coffea canephora* is an economically important species, found in the understorey of tropical rainforests of Central and West Africa, and the genetic diversity harboured in its wild populations is vital for sustainable coffee production worldwide. Here, we aimed to quantify genetic diversity, genetic structure, and pedigree relations in wild *C. canephora* populations, and we investigated associations between these descriptors and forest disturbance and regeneration. Therefore, we sampled 256 *C. canephora* individuals within 24 plots across three forest categories in Yangambi (DR Congo), and used genotyping-by-sequencing to identify 18 894 SNPs. Overall, we found high genetic diversity, and no evidence of genetic erosion in *C. canephora* in disturbed old-growth forest, as compared to undisturbed old-growth forest. Additionally, an overall heterozygosity excess was found in all populations, which was expected for a self-incompatible species. Genetic structure was mainly a result of isolation-by-distance, reflecting geographical location, with low to moderate relatedness at finer scales. Populations in regrowth forest had lower allelic richness than populations in old-growth forest and were characterised by a lower inter-individual relatedness and a lack of isolation-by-distance, suggesting that they originated from different neighbouring populations and were subject to founder effects. Wild Robusta coffee populations in the study area still harbour high levels of genetic diversity, yet careful monitoring of their response to ongoing forest degradation remains required.

## Introduction

Tropical rainforests cover only 7% of the Earth’s land surface but represent the world’s richest reservoir of terrestrial biodiversity (Kier et al. 2005; Kreft & Jetz 2007). Over the past decades, human activities such as industrial logging and the encroachment of agriculture and infrastructure have negatively impacted tropical forest cover and resulted in the loss of biodiversity, jeopardising the provisioning of important ecosystem services such as carbon sequestration and climate regulation (Gardner et al. 2009; Curtis et al. 2018; Edwards et al. 2019). Less conspicuous than the loss of tropical forest cover is the ongoing degradation of tropical forests. Forest degradation refers to within-forest disturbance under a more or less intact canopy, and is mainly caused by selective logging and the removal of the understorey vegetation (Sasaki & Putz 2009; Tyukavina et al. 2018). Degradation of tropical forests may be as detrimental to biodiversity as forest cover loss, due to the large spatial scales at which it occurs (Barlow et al. 2016). In the Congo Basin, for example, the rate of forest degradation has been estimated at 317,000 ha per year between 2000 and 2005 (Ernst et al. 2013), whereas Shapiro et al. (2021) reported that 23 million ha of forest has been degraded between 2000 and 2016 in this region.

Forest degradation may compromise the resilience and long-term stability of tropical rainforests because it can negatively affect the regeneration of the remaining woody plant species (Norden et al. 2009). Plant regeneration and fitness depend on multiple processes, including pollination, seed dispersal, germination and seedling establishment (Barrett & Eckert 1990). Crucial aspects of gene flow early in the regeneration cycle, such as pollination and seed dispersal, can become strongly jeopardised through ongoing large-scale anthropogenic disturbance of tropical forests (Neuschulz et al. 2016). Because many tropical canopy trees and understorey shrubs typically occur in population densities of less than one individual per ha, and due to widespread dioecy and self-incompatibility (SI) (Bawa et al. 1985; Hubbell & Foster 1986), pollen flow and successful pollination and reproduction can be expected to be particularly susceptible to changes in the understorey plant species density and composition (Aguilar et al. 2019; Chiriboga-Arroyo et al. 2021). Reduced gene flow may not only result in decreased reproductive capacity, but also in changes in the genetic structure and in genetic erosion of the remaining populations through increased genetic drift and inbreeding (Vranckx et al. 2012; Ismail et al. 2017; Campbell et al. 2018). This process can be exacerbated by the disappearance of large frugivores (by hunting, or as a result of habitat loss) from disturbed tropical rainforests (Bello et al. 2015), hampering seed dispersal and recruitment. Ultimately, reduced pollen flow and seed dispersal may even result in the local extinction of shrub and tree species (da Silva & Tabarelli 2000).

Apart from tropical forest disturbance, tropical forest regeneration on abandoned agricultural land may also significantly impact genetic diversity and structure of the recolonising woody species. Such regrowth forests make up an increasing fraction of the forested area throughout the tropics (FAO & UNEP 2020; Poorter et al. 2021). Recolonisation of abandoned agricultural fields by tropical woody species almost entirely depends on seed dispersal, as these species usually do not have a persistent soil seedbank (Sezen et al. 2007). These colonisation events are expected to be prone to founder effects, in which the newly founded population represents only a subsample from one or a few neighbouring source populations (Wright 1932; Mayr 1954; Widmer & Lexer 2001). These founder effects can result in major genetic changes, including loss of genetic diversity and increased genetic differentiation among populations, with associated fitness consequences (Born et al. 2008; Vandepitte et al. 2012). Whereas ample research has already been done on species diversity and community composition in regrowth tropical forests (e.g., Oberleitner et al. 2021; Makelele et al. 2021; Depecker et al. 2022), studies on the genetic diversity of tropical tree species in regrowth forests in the tropics are still scarce, especially in Africa.

*Coffea canephora* (Robusta coffee) is an understorey tree from the lowland tropical rainforests of Central and West Africa. The conservation of its genetic diversity is of utmost importance for future sustainable coffee production worldwide as wild populations carry useful traits for coffee breeding, such as disease resistance (Silva et al. 2006; Lashermes et al. 2010), tolerance to climate change (Davis et al. 2012) and drought tolerance (Cramer 1957). Robusta coffee currently accounts for more than 40% of the global coffee production (ICO 2022) but is gaining commercial importance thanks to its higher disease resistance (Leroy et al. 2005), higher productivity (Wellman 1961) and its assumedly lower susceptibility to climate change than Arabica coffee (Craparo et al. 2015; Davis et al. 2012). *Coffea canephora* is a self-incompatible species, without a persistent soil seed bank (Oryem-Origa 1999; Nowak et al. 2011). Natural populations of *C. canephora* are usually disconnected, with 10 to 20 individuals per ha, and few offspring scattered across the forest floor (Musoli et al. 2009; Cubry et al. 2013; Depecker & Vandelook pers. obs.). Such characteristics can be expected to render the genetic diversity and structure of this species very susceptible to both the processes of forest disturbance and forest regrowth. Yet, research on the genetic diversity of wild *C. canephora* is still rare (but see Musoli et al. 2009; Kiwuka et al. 2021; Vanden Abeele et al. 2021 at the nationwide scale; and Nyakaana 2007 at the population scale).

In this study, we aimed to quantify the association between rainforest disturbance and regrowth on the one side, and genetic diversity and genetic structure of wild *C. canephora* on the other side, focusing on the Yangambi area (DR Congo) in the Congo Basin, an important Robusta coffee genetic diversity hotspot (Ferrao et al. 2019; Merot-l’Anthoene et al. 2019). Therefore, we surveyed 24 inventory plots across undisturbed old-growth forest, disturbed old-growth forest, and regrowth forest, in which a total of 256 *C. canephora* individuals were sampled, and genotyped using genotyping-by-sequencing (GBS). We hypothesised to find (i) lower genetic diversity in disturbed old-growth forest and regrowth forest, as compared to undisturbed old-growth forest, (ii) more pronounced genetic structure and pedigree relations in disturbed old-growth forest, and (iii) that populations in regrowth forests have emerged through colonisation via seed dispersal from multiple neighbouring coffee populations in old-growth forest, resulting in strongly admixed populations.

## Material and methods

### Study population and sampling

The Yangambi region is located in the Tshopo province in North-Eastern DR Congo, approximately 100 km west of Kisangani. The Yangambi landscape consists of a mosaic of land tenures, typical for the Congo Basin: the Yangambi Man and Biosphere Reserve; the Ngazi Forest Reserve; a logging concession; and customary land (van Vliet et al. 2018).

Previously, Depecker et al. (2022) established 25 forest inventory plots of 125 m × 125 m (1.56 ha), covering an area of ca. 50-by-20 km, just North of the Congo River (**Fig. 1A**). We adopted their classification of the plots into three different forest categories: (i) Plots in regrowth forest (8 plots) located on historical agricultural land. Depecker et al. (2022) estimated that these agricultural lands were abandoned somewhere between 1962 and 1980, and since then overgrown. (ii) Plots in disturbed old-growth forest (7 plots), with clear indications of small-scale selective logging through the presence of tree stumps and (iii) plots in undisturbed old-growth forest (10 plots) without signs of disturbance.

**Figure 1:**
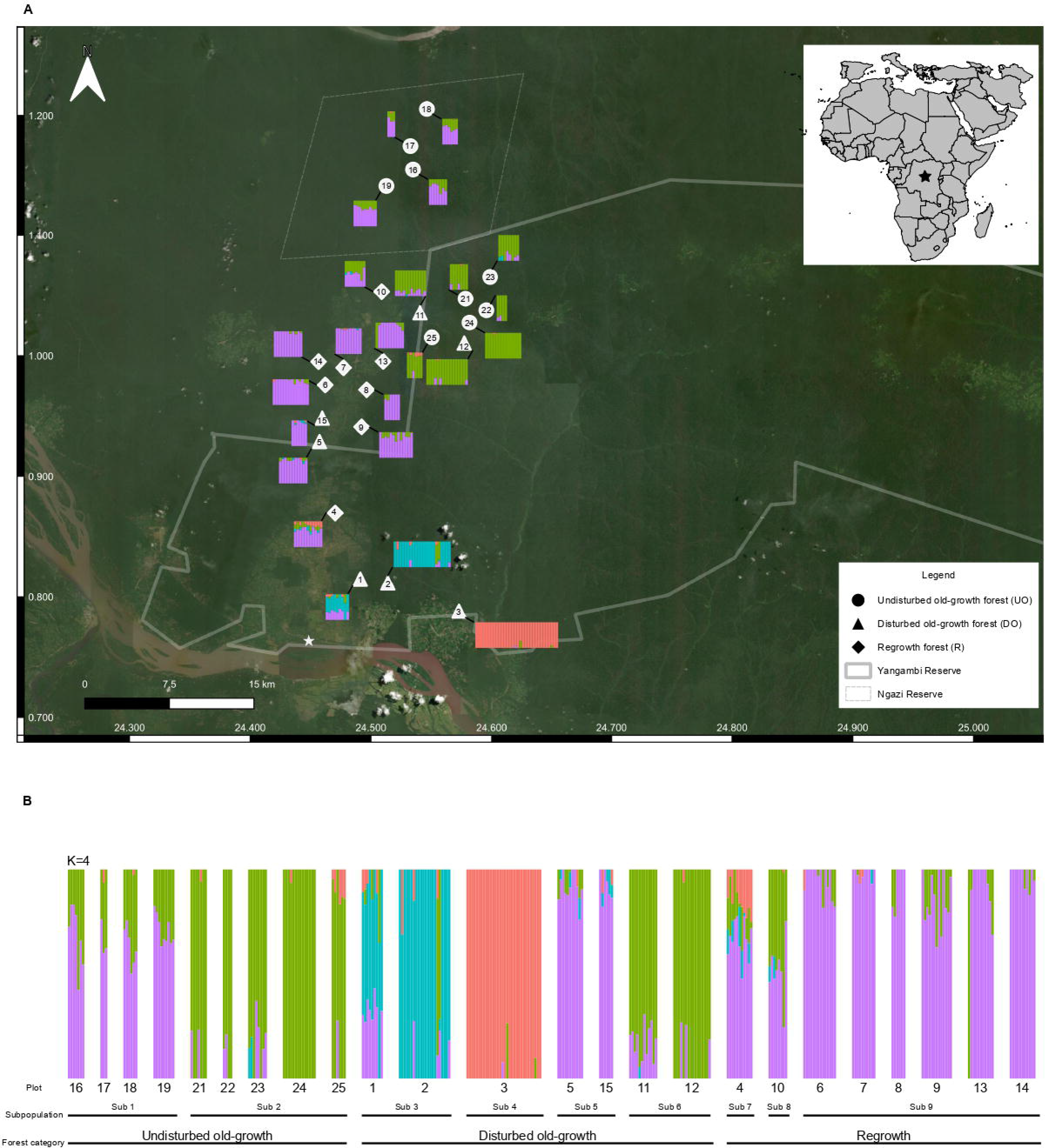
Estimation of subpopulations using 256 wild *C. canephora* individuals and 794 SNPs. (A) Map of the Yangambi region showing the location of each plot as well as the K coloured segments (K=4) of each individual within each plot. The white star on the map locates the commune of Yangambi. (B) The entire population of *C. canephora* was divided into 4 clusters (K=4) using fastSTRUCTURE. Individuals are shown by thin vertical lines, which are divided into K coloured segments representing the estimated membership probabilities (Q) of each individual.

In this study, these plots were systematically surveyed for *C. canephora* by multiple Afrotropical plant experts. A leaf sample of all *C. canephora* individuals was collected and silica-dried, yielding a total of 256 samples. One survey plot (plot #20) from Depecker et al. (2022) was omitted, because there were too few *C. canephora* individuals to adequately analyse.

### Genomic DNA extraction and Genotyping-by-Sequencing (GBS)

20-30 mg leaf material was homogenised with a Retsch TissueLyser II (Mixer Mill MM 500 Nano; Retsch^®^). Genomic DNA was extracted from the dried leaf material using an optimised cetyltrimethylammonium bromide (CTAB) protocol adapted from Doyle & Doyle (1987). DNA quantities were measured with the Quantifluor dsDNA system on a Promega Quantus Fluorometer (Promega, Madison, USA).

GBS libraries were prepared using a double-enzyme GBS protocol adapted from Elshire (2011) and Poland et al. (2012). In short, 100 ng of genomic DNA was digested with *PstI* and *MseI* restriction enzymes (New England Biolabs, Ipswich, USA), and barcoded and common adapter constructs were ligated with T4 ligase (New England Biolabs, Ipswich, USA) in a final volume of 35 μL. Ligation products were purified with 1.6x MagNA magnetic beads (GE Healthcare Europe, Machelen, BE) and eluted in 30 μl TE. Of the purified DNA eluate, 3 μl was used for amplification with *Taq* 2x Master Mix (New England Biolabs, Ipswitch, USA) using a 18 cycles PCR protocol. PCR products were bead-purified with 1.6x MagNA, and their DNA concentrations were quantified using a Quantus Fluorometer. The library quality and fragment size distributions were assessed using a QIAxcel system (Qiagen, Venlo, NL). Equimolar amounts of the GBS libraries were pooled, bead-purified and 150 bp paired-end sequenced on an Illumina HiSeq-X instrument by Admera Health (South Plainfield, USA).

### Data processing

Reads were processed with a customised script available on Gitlab (https://gitlab.com/ilvo/GBprocesS). The quality of sequence data was validated with FastQC 0.11 (Andrews 2010) and reads were demultiplexed using Cutadapt 2.10 (Martin 2011), allowing zero mismatches in barcodes or barcode-restriction site remnant combination. The 3’ restriction site remnant and the common adapter sequence of forward reads and the 3’ restriction site remnant, the barcode, and the barcode adapter sequence of reverse reads were removed based on sequence-specific pattern recognition and positional trimming using Cutadapt. After trimming the 5’ restriction site remnant of forward and reverse reads using positional trimming in Cutadapt, forward and reverse reads with a minimum read length of 60 bp and a minimum overlap of 10 bp were merged using PEAR 0.9.11 (Zhang et al. 2014). Merged reads with a mean base quality below 25 or with more than 5% of the nucleotides uncalled and reads containing internal restriction sites were discarded using GBprocesS. Merged reads were aligned to the *C. canephora* reference genome sequence (Denoeud et al. 2014) with the BWA-mem algorithm in BWA 0.7.17 with default parameters. Alignments were sorted, indexed, and filtered on mapping quality above 20 with SAMtools 1.10 (Li et al. 2009).

Single nucleotide polymorphisms (SNPs) were called with GATK (Genome Analysis Toolkit) Unified Genotyper 3.7.0. (McKenna et al. 2010). Multi-allelic SNPs were removed with GATK and the remaining SNPs were filtered using the following parameters: min-meanDP 30, mac 4, minQ 20 and minimal 80% completeness. The remaining SNPs were then subjected to further filtering with the following parameters: minDP10, minGQ 30, max-missing 0.7, mac 3, minQ 30, min-alleles 2, max-alleles 2 and maf 0.05 using VCFtools 0.1.16 (Danecek et al. 2011).

### Genetic diversity and structure analysis

The number of effective alleles (N_e_) was calculated in GenAlEx 6.5 (Peakall & Smouse 2012) for each plot and forest category. Allelic richness (A_r_) was calculated according to El Mousadik and Petit (1996) using the *allelic.richness* function of R package *hierfstat* (Goudet 2013). The observed and expected heterozygosity (H_o_ and H_e_, respectively) and inbreeding coefficient (F_IS_) were calculated in VCFtools, for each plot and forest category. All genetic diversity indices were compared between the three forest categories using non-parametric Kruskal-Wallis rank sum tests, followed by Dunn’s Multiple Comparison tests. To estimate the genetic distance between all coffee plots, pairwise F_ST_ values (Weir & Cockerham 1984) were calculated using PLINK 1.9 (Chang 2015). Mantel tests (Podani 2000) were performed in RStudio to test the correlation between genetic (F_ST_) and geographical distances across all plots, and across plots within each of the three forest categories.

To comply with the assumptions for a genetic structure analysis, SNPs were filtered, using VCFtools based on Hardy-Weinberg Equilibrium (hwe 0.01), minor allele frequencies (maf 0.05) and linkage disequilibrium (indep-pairwise 50 10 0.5). Discriminant analysis of principal components (DAPC) (Jombart & Collins 2015) was then conducted using the R package ADEGENET (Jombart 2008; RStudio Team 2016). First, the *find.clusters* function, which runs successive K-means clustering with increasing number of clusters (k), was used to assess the number of clusters that maximises *between*-group variance and minimises *within*-group variance. The Bayesian Information Criterion (BIC) was applied to select the most optimal value of k. DAPC was then performed on the most optimal number of clusters (k) using the *dapc* function. Second, a Bayesian clustering implemented in fastSTRUCTURE 1.0 (Raj et al. 2014) was run to assess genetic structure in the *C. canephora* individuals given the most optimal number of genetic clusters (K). Hundred iterations were run for each expected cluster setting K, ranging from 2 to 9. The StructureSelector software (Li & Liu 2018) was used to determine the most optimal number of K, by first plotting the mean log probability of each successive K and then using the Delta K method following Evanno et al. (2005). Graphical representation of the fastSTRUCTURE results was done in RStudio.

### Relatedness and relationships

To reveal patterns of historical gene flow, we analysed the relatedness and relationships among the 256 individuals using multi-allelic short haplotypes (created via read-backed phasing of the SNPs with the SMAP package) which were preferred over single bi-allelic SNPs because of their increased information content (Schaumont et al. 2022). Furthermore, the use of haplotypes reduces the optimum number of SNPs needed to achieve high assignment success rates in complex scenarios (Garcia-Fernandez et al. 2018). Read-backed haplotyping of the SNPs was done with SMAP haplotype-sites using the optimal parameter settings for diploid individuals and double-enzyme GBS merged reads with: no-indels, min-haplotype-frequency 5, discrete calls dosage, dosage-filter 2, min-read-count 10, min-distinct-haplotypes 2, max-distinct-haplotypes 10 frequency-interval-bounds default for diploids, and locus-correctness 90%. Subsequently, SMAPapp-Matrix (https://gitlab.com/ybawin/smapapps) was used to calculate the locus information content (LIC). The criteria were set so that all loci with at least one unique haplotype were considered. Based on this criterion, we selected the haploset with the 100 most informative loci. The strength of the LIC is that informative loci with more than two haplotypes are retained even if the minor haplotype frequency is low, and thus LIC is a better criterion for the discriminatory power of all haplotypes per locus.

The relatedness (r), based on maximum likelihood, was calculated among all pairs of individuals using ML-Relate (Kalinowski et al. 2006). The relatedness between individual pairs represents the overall identity-by-descent in a continuous measure, ranging between zero and one (Blouin 2003). Mean relatedness was calculated afterwards between pairs of individuals at both the plot-level and the forest category-level. To evaluate differences between forest categories, a Kruskal-Wallis rank sum test and Dunn’s Multiple Comparisons test was used.

In addition, and also using ML-Relate, pairs of individuals were classified into four pedigree relations using maximum likelihood estimates: unrelated (U), half-siblings (HS), full-siblings (FS), and parent-offspring (PO) (Jones et al 2010). Log-likelihoods were calculated for all four relationships and the one with the highest value was assigned to the corresponding tested pair of individuals. Afterwards, the frequency of each assigned relationship was counted at both plot-level and forest category-level. Differences in frequencies of relationships at the category-level were tested using a Pearson’s chi-squared test and pairwise Pearson’s chi-squared tests with simulated p-values based on 9999 replicates, using the chisq.test function in the *stats* package (RStudio Team 2016).

## Results

### SNP discovery and selection

A total of 18 894 biallelic SNPs with a completeness of at least 80% were identified across all individuals in undisturbed old-growth forest (n=64 coffee samples), disturbed old-growth forest (n=108), and regrowth forest (n=84) plots. Of these, 3 212 SNPs with a minimum minor allele count of 3 and a minimum minor allele frequency of 0.05 were used for the genetic diversity analysis. Yet, we used only 794 SNPs for the genetic structure analysis, which had been filtered based on the Hardy-Weinberg and linkage disequilibrium criteria.

### Genetic diversity

Genetic diversity measures for all plots and forest categories are presented in **Table 1**. The number of effective alleles (N_e_) was significantly lower in plots in undisturbed old-growth forest than in plots in disturbed old-growth forest (p=0.006), but N_e_ in plots in regrowth forest was not significantly different from plots in disturbed and undisturbed old-growth forest (p=0.10 and p=0.11, respectively). Allelic richness (A_r_) was significantly lower in plots in undisturbed old-growth forest compared to plots in disturbed old-growth forest (p<0.001) and significantly lower in plots in regrowth forest compared to both plots in disturbed and undisturbed old-growth forest (p=0.03 and p<0.001, respectively). No significant differences were found in observed heterozygosity (H_o_), expected heterozygosity (H_e_) and inbreeding coefficient (F_IS_) between plots from the three forest categories (p=0.16, p=0.85, p=0.15, respectively).

**Table 1:**
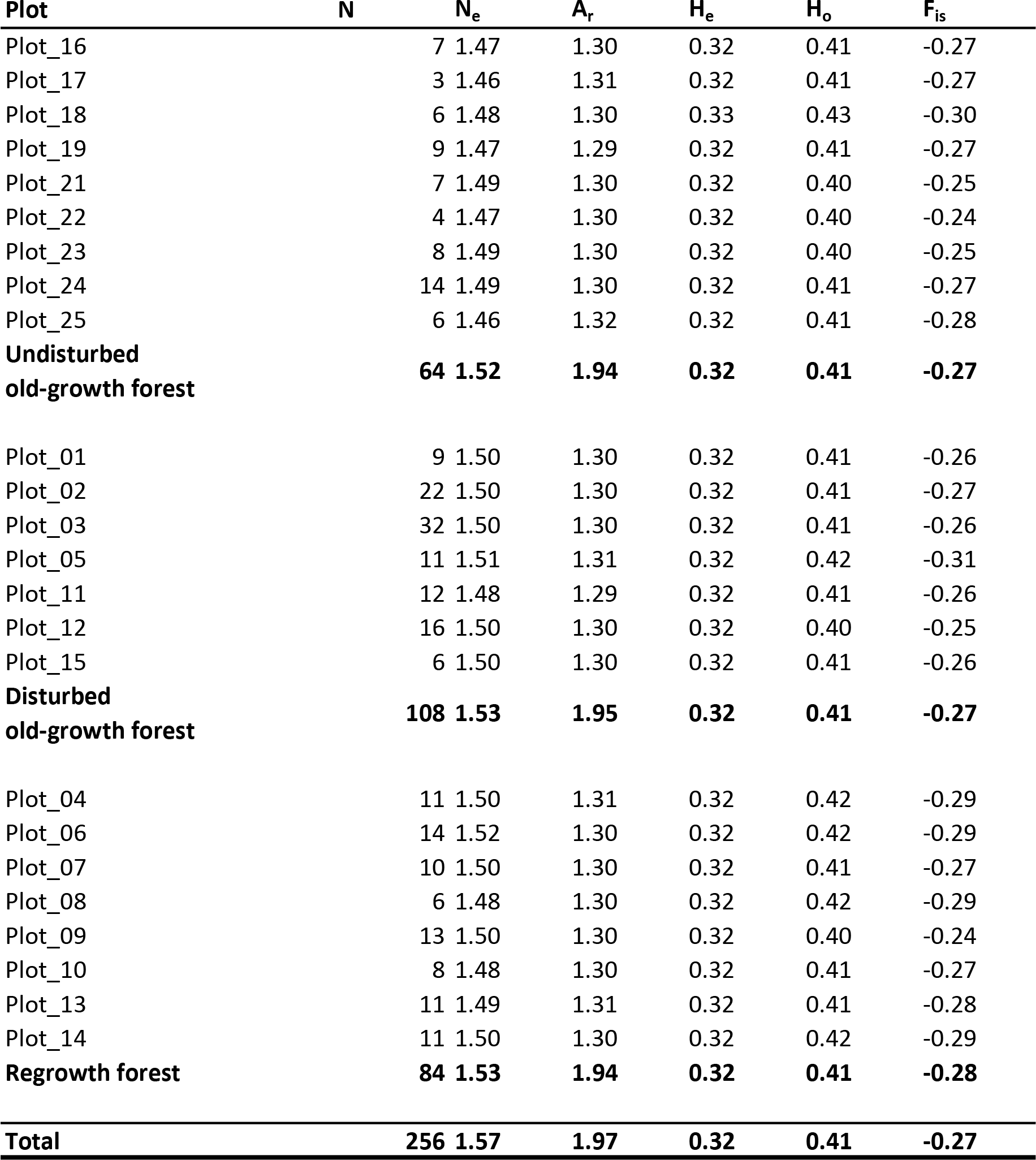
Genetic diversity estimates for all *C. canephora* sampling plots across the three forest categories in the Yangambi region in DRC. N = sample size; N_e_= Effective number of alleles, A_r_= allelic richness; H_o_= Observed heterozygosity; H_e_ = Expected heterozygosity, F_IS_= inbreeding coefficient.

Pairwise genetic differentiation (F_ST_) was highest among plots in disturbed and plots in undisturbed old-growth forest (F_ST_=0.025). Genetic differentiation was higher among plots in undisturbed old-growth forest and plots in regrowth forest (F_ST_=0.025) than among plots in disturbed old-growth forest and plots in regrowth forest (F_ST_=0.017). Isolation-by-distance was found across all plots (Mantel r-statistic=0.24, p=0.01; **Fig. 2A**) and across the plots within disturbed and undisturbed old-growth forest separately (undisturbed forest: Mantel r-statistic=0.48, p=0.024, **Fig. 2B**; disturbed forest: Mantel r-statistic=0.47, p=0.036, **Fig. 2C**). No significant isolation-by-distance was found across plots in regrowth forest (Mantel r-statistic=0.24, p=0.2; **Fig. 2D**).

**Figure 2:**
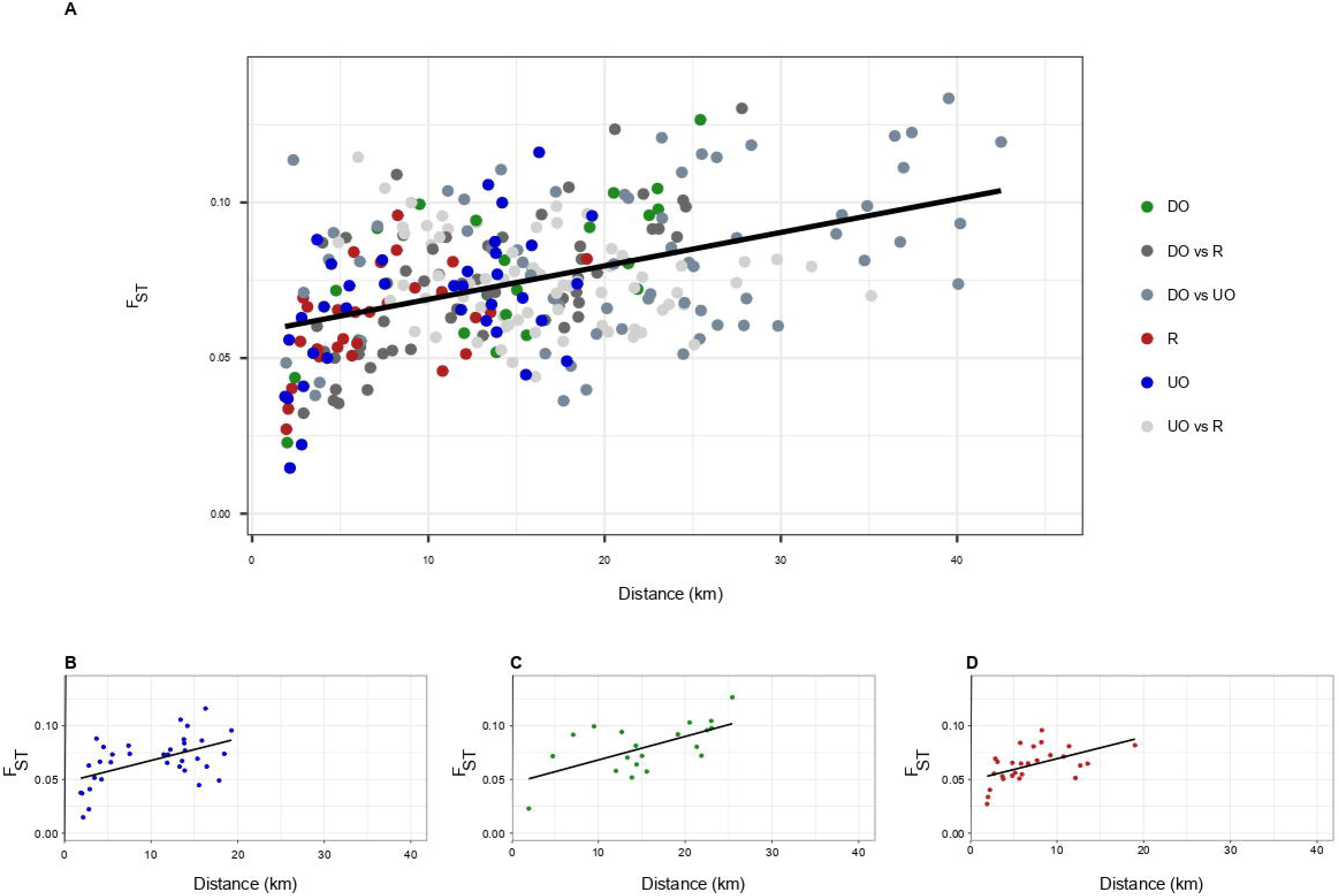
Relationship between geographic (kilometres) and genetic (F_ST_) distances between *C. canephora* populations in different forest categories of the Yangambi region. The lines show the positive relationship between both variables. Relationship shown over (A) the entire population of *C. canephora*; (B) all individuals located in undisturbed old-growth forest (UO, blue); (C) all individuals located in disturbed old-growth forest (DO, green); (D) all individuals located in regrowth forest (R, red).

### Genetic structure analysis

Four different clusters were identified using the DAPC analysis, performed on the first hundred PCs of the PCA and three discriminant eigenvalues. Cluster 1, containing samples from plot 3, was separated from the other clusters according to Linear Discriminant 1 (LD1) (Supplementary, **Fig. S1B**).

The fastSTRUCTURE analysis showed that the plots in undisturbed old-growth forest were divided into two subpopulations (**Fig. 1B**), namely plot 16 to 19 and plot 21 to 25, with individuals in plots 16 to 19 showing a mixture of cluster 2 and 4 (**Fig. 1A**). Plots in disturbed old-growth forest were divided into four subpopulations (**Fig. 1B**), namely: plots 1 and 2; plot 3; plots 5 and 15; plots 11 and 12 (**Fig. 1A**). Plots in regrowth forest were divided into three subpopulations (**Fig. 1A**), namely: plot 4; plot 10; plots 6 to 9, 13 and 14. Individuals in plot 10 showed to be a mixture of clusters 2 and 4, whereas individuals in plot 4 showed a mixture of all clusters. Overall, clustering of the old-growth samples was consistently differentiated according to the geographical location of the plots (**Fig. 1A**).

### Relatedness and relationships

The average relatedness values per plot ranged between 0.074 and 0.304, with an average value of 0.184. The relatedness between pairs of individuals was significantly different among forest categories (χ^2^=65.27, p<0.001) (**Fig. 3A**). Specifically, it was higher in undisturbed old-growth forest than in regrowth forest (p<0.001) and higher in disturbed old-growth forest than in regrowth forest (p<0.001). There were no significant differences between undisturbed old-growth forest and disturbed old-growth forest.

**Figure 3:**
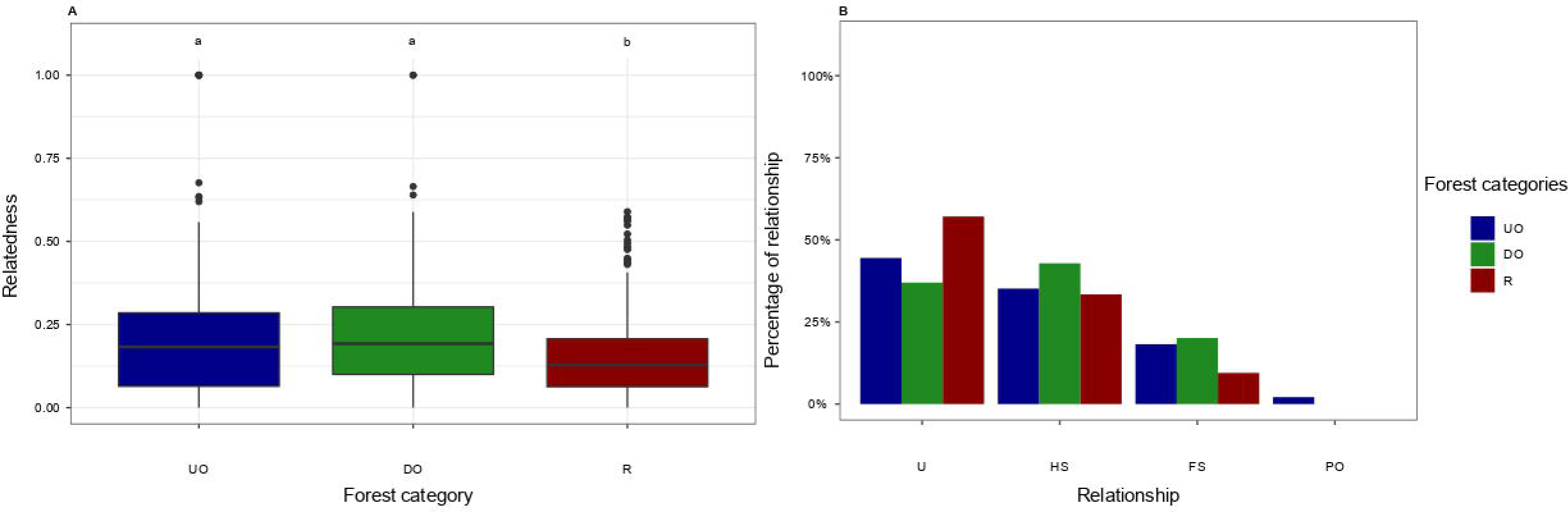
Determination of genetic relatedness and relationships between different forest categories. A) Estimated relatedness between undisturbed old-growth forest (UO), disturbed old-growth forest (DO) and regrowth forest (R). B) Estimated frequencies of unrelated individuals, half-siblings, full-siblings and parent-offspring relationships within undisturbed old-growth (UO), disturbed old-growth (DO) and regrowth (R) forest. Letters code for significantly different groups.

The frequency distribution of the relationship (unrelated, half-siblings, full-siblings, parent-offspring) of pairs of individuals within plots was significantly different among forest categories (χ^2^=86.543, p<0.05). Specifically, significant differences in the frequency distribution were found among plots in undisturbed old-growth and disturbed old-growth forest (χ^2^=27.49, p<0.001), plots in undisturbed old-growth forest and regrowth forest (χ^2^=22.83, p<0.001), and plots in disturbed old-growth forest and regrowth forest (χ^2^=54.69, p<0.001), (**Fig. 3B**). Parent-offspring pairs were only found in undisturbed old-growth forest, and even then, only at low incidence, namely: two pairs in plot 21; one in plot 22; one in plot 23; one in plot 24.

## Discussion

Understanding the genetic variation and its relation with anthropogenic disturbance is of major importance for the conservation of *C. canephora* genetic resources in the Congo Basin. Our study encompasses the most densely sampled set of wild individuals of *C. canephora* so far. By using GBS-derived SNP markers, we were able to quantify genetic diversity, map genetic structure, and determine pedigree relations in *C. canephora* populations and compare these indicators among undisturbed old-growth forest, disturbed old-growth forest, and regrowth forest.

## Genetic diversity

A high genetic diversity, both in terms of allelic diversity and heterozygosity, was found in all 24 sampling plots across the three forest categories. Our findings are in line with other studies that have used SSR markers, such as Vanden Abeele et al. (2021), who found high heterozygosity (H_o_=0.48) in wild *C. canephora* populations in the Tshopo Province of DR Congo (Yangambi and Yoko, Kisangani). Likewise, Nyakaana (2007) found a high mean observed heterozygosity (H_o_=0.46) across five localities in the Kibale National Park in Uganda. Elsewhere in Uganda, Musoli et al. (2009) found a mean H_o_ of 0.37 over two separate regions, while Kiwuka et al. (2021) found a mean H_o_ of 0.51 over seven distinct regions. All our sampling plots harboured a higher H_o_ than reported over the whole Guineo-Congolian region, where H_o_ ranged between 0.27 and 0.38 (Gomez et al. 2009; Cubry et al. 2013). This suggests that the Yangambi area is key for the conservation of *C. canephora* genetic resources.

We hypothesised that anthropogenic disturbance leads to decreased population genetic diversity, specifically due to selective logging (Depecker et al. 2022). However, we found no evidence of reduced genetic diversity in plots in disturbed old-growth forest, as compared to plots in undisturbed old-growth forest. On the contrary, the number of effective alleles and allelic richness were significantly lower in plots in undisturbed old-growth forest, compared to plots in disturbed old-growth forest. One explanation could be historical spatial variation in genetic diversity. Unfortunately, there is no information on the historical genetic diversity of wild *C. canephora* in the Yangambi area. An alternative explanation for the higher genetic diversity could be increased levels of gene flow in disturbed areas, contrary to our expectations. Several studies have found enhanced pollinator activity through disturbance, promoting gene flow (Dick et al. 2003 and references therein). It is possible that forest disturbance altered the pollinating insect communities, with for instance a higher abundance of *Apis mellifera*, which can govern long-distance pollination as has been observed in the tropical Amazonian tree species *Dinizia excelsa* (Dick et al. 2003). Furthermore, competition for space and resources is likely to be reduced in disturbed areas, possibly resulting in higher seed survival and germination rates, which can lead to better seedling establishment (Olsson et al. 2019).

In addition, we found that, in general, observed heterozygosity was higher than expected heterozygosity, indicating an excess of heterozygotes. Such negative values of F_IS_ are in accordance with obligate outcrossing in self-incompatible plant species like *C. canephora* (Mateu-Andrés & De Paco 2006).

## Genetic structure

Limited genetic connectivity between plots may have resulted in relatively high genetic differentiation among our sampled plots at relatively short geographical distance. Likewise, in the tropical rainforest of West Uganda, Nyakaana (2007) detected strong genetic differentiation between five *C. canephora* populations, which were only separated by short geographical distances. Similar patterns were detected in other tropical woody plant species, including several *Psychotria* species (Theim et al. 2014), *Paypayrola blanchetiana* (Braun et al. 2020), and *Theobroma cacao* (Lachenaud et al. 2008).

The genetic differentiation was also reflected in the significantly genetically diverged clusters, which were demonstrated by DAPC analysis and further supported by the fastSTRUCTURE analysis. It is remarkable that the *C. canephora* individuals sampled in plot 3 were markedly genetically separated from the individuals sampled in all other plots. This may be explained by the monodominant and species-poor *Gilbertiodendron dewevrei* forest that forms a natural barrier and isolates plot 3 from the other ones (Kearsley et al. 2017). In general, this type of monodominant forests significantly alters the understorey environment, making it difficult for other species to establish and survive (Torti et al. 2001). *Coffea canephora* has never been observed in the *G. dewevrei* forest understorey (Asimonyio & Kambale, pers. obs.), but more sampling in the vicinity of plot 3 is needed to confirm this hypothesis.

Geographic location, rather than anthropogenic disturbance, appears to be the main driver of genetic structure across the whole study area. The identified subclusters can be attributed to two different gradients in terms of geographic distance, and which are clearly subjected to isolation-by-distance. Firstly, a more or less east-west gradient separating plots 11 and 21 to 25 from the other plots. Secondly, a north-south gradient, visible in the more continuously sampled area. Within this north-south gradient, plots in the south are clearly more differentiated and admixed than plots in the north. Within the Yangambi region, anthropogenic activity is high close to the Congo River. Nevertheless, further research is necessary to test the association between the anthropogenic activities and the higher rate of differentiation in plots in the south. Plots in regrowth forest follow the same patterns of gene flow and show high levels of admixture but are not subjected to isolation-by-distance. These findings suggest that after agricultural abandonment, *C. canephora* individuals coming from neighbouring old-growth forests recolonised this area. This is further supported by the lower allelic diversity, which hints at founder effects, possibly due to a limited number of migrants.

## Pedigree relations

By assessing the relatedness and relationships between pairs of individuals, we were able to reveal putative patterns of dispersal and recruitment. Overall, we found a low to moderate relatedness among pairs of individuals within one plot. Furthermore, most pairs of individuals in the majority of the plots were unrelated or half-siblings. Combined with the observed genetic structure, we hypothesise that gene flow is limited at larger distances (between plots), as also indicated by the significant isolation-by-distance and the observation that no parent-offspring pairs were found between plots, with a minimal distance of two km. This distance-dependent decay of gene flow is consistent with the theory of isolation-by-distance models (Vekemans & Hardy 2004).

A lower relatedness was detected within plots in regrowth forest as compared to within plots in undisturbed old-growth forest and in disturbed old-growth forest. This confirms that these regrowth areas were colonised by *C. canephora* migrants from multiple neighbouring sources, thus, lowering the average relatedness among individuals within plots in regrowth forests. This pattern is supported by the common observation that relatedness decreases as migration increases (Jones & Wang 2012).

## Conclusion

We found that the wild *C. canephora* populations in the Yangambi region harbour both a high allelic diversity and heterozygosity, thereby pointing at the importance of the wild *C. canephora* populations in the Congo Basin as hotspots of genetic diversity. Because local studies on the genetic diversity of wild *C. canephora*, and by extension other rainforest understorey species, are very rare in the Congo Basin, our study can be used as a reference for future research, in which novel quantifications of genetic diversity can be compared with the values found in our work. Indeed, although we could not detect genetic erosion in disturbed forests, it is important to continue monitoring the effect of anthropogenic disturbance on the genetic diversity, genetic structure, and gene flow in wild populations of *C. canephora*, because the observations made in this study might be influenced by the historical distribution of genetic diversity. Conservation of the genetic diversity and actors governing gene flow in old-growth forests is crucial, although the populations in regrowth forests can aid in the maintenance of the genetic resources, which are important for the future of coffee cultivation.

## Acknowledgments

We would like to thank the Institut National pour l’Étude et la Recherche Agronomiques (INERA) and the FORETS project, which is financed by the 11^th^ European Development Fund, for facilitating the field mission. We would also like to express our sincere gratitude to the Ministère de L’Environnement et Développement Durable (MEDD) for their help with obtaining permits (N°008/ANCCB-RDC/SG-EDD/BTB/11/2020 & N°001/ANCCB-RDC/SG-EDD/BTB/01/2021).

## Author contributions

OH, FV, TR, JD and LV designed this study. JD, JA, YH, JK, IMM, TE participated in fieldwork. LV and AS executed the lab work. JD, LV and YB analysed the data. JD, LV, FV, OH, TR wrote the manuscript. All authors contributed to finalising the manuscript.

## Data availability statement

FASTQ read files of all GBS libraries were deposited at the NCBI sequence read archive (SRA) in BioProject (projectnummer will be added).

## Supplemental information

Supplemental Figure 1: A) Number of *C. canephora* individuals for each plot divided over each genetic cluster. B) DAPC of genetic clusters present in entire population of *C. canephora*. The inset shows number of PCs that were retained from the PCA analysis (lower left corner) and the Linear Discriminants retained from the discriminate analysis (upper left corner).

Supplemental Figure 2: Determination of genetic relatedness and relationships for each plot. A) Estimated relatedness for each plot within undisturbed old-growth (UO), disturbed old-growth (DO) and regrowth (R) forest. B) Estimated frequencies of unrelated individuals, half siblings, full siblings and parent-offspring relationships for each plot within undisturbed old-growth (UO), disturbed old-growth (DO) and regrowth (R) forest. Letters code for significantly different groups.

