## Supplementary figures and images for "Genetic diversity and structure in wild Robusta coffee (*Coffea canephora* A. Froehner) populations in Yangambi (DR Congo) and their relation with forest disturbance"

### Supplemental figure 1

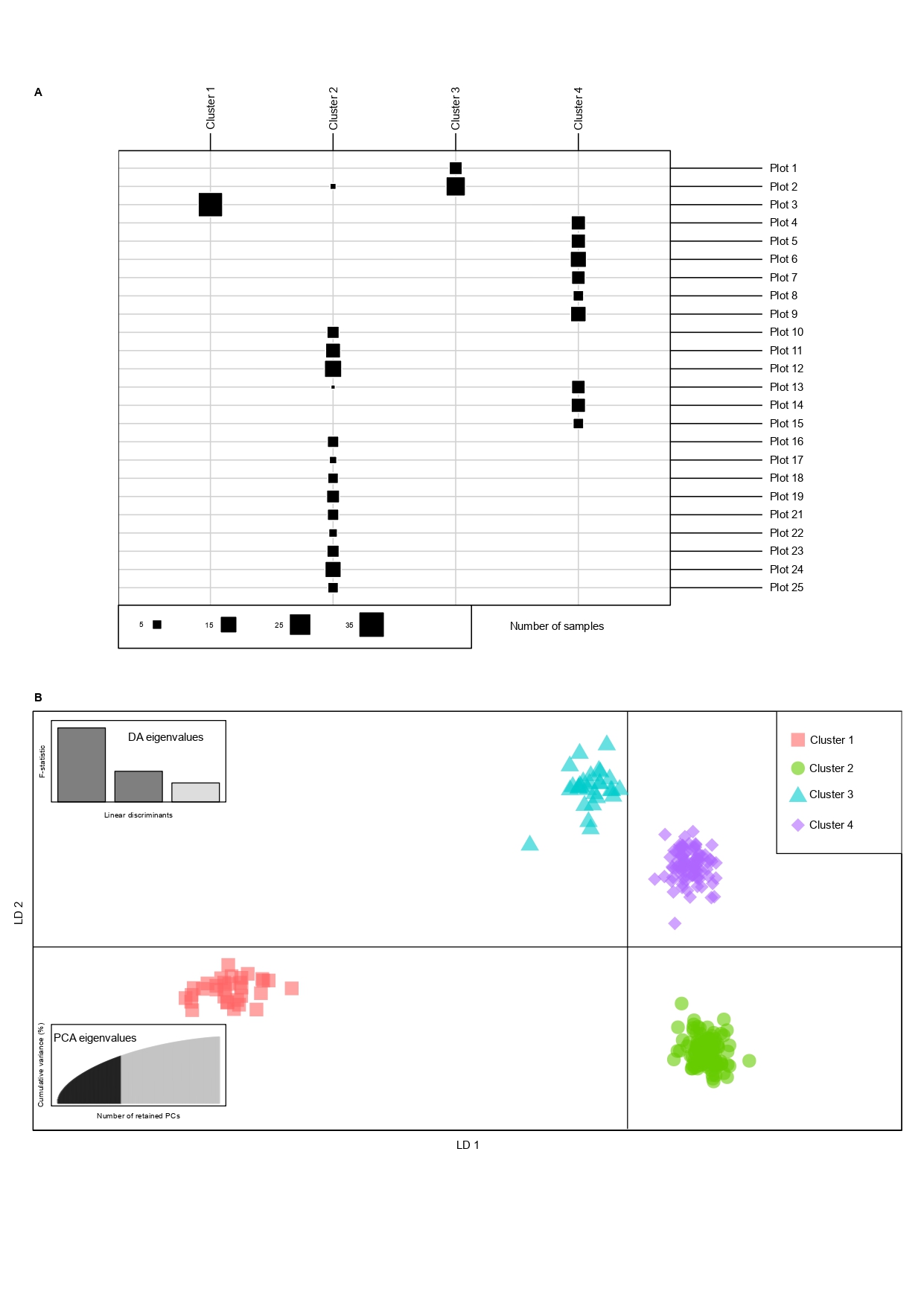

### Supplemental figure 2

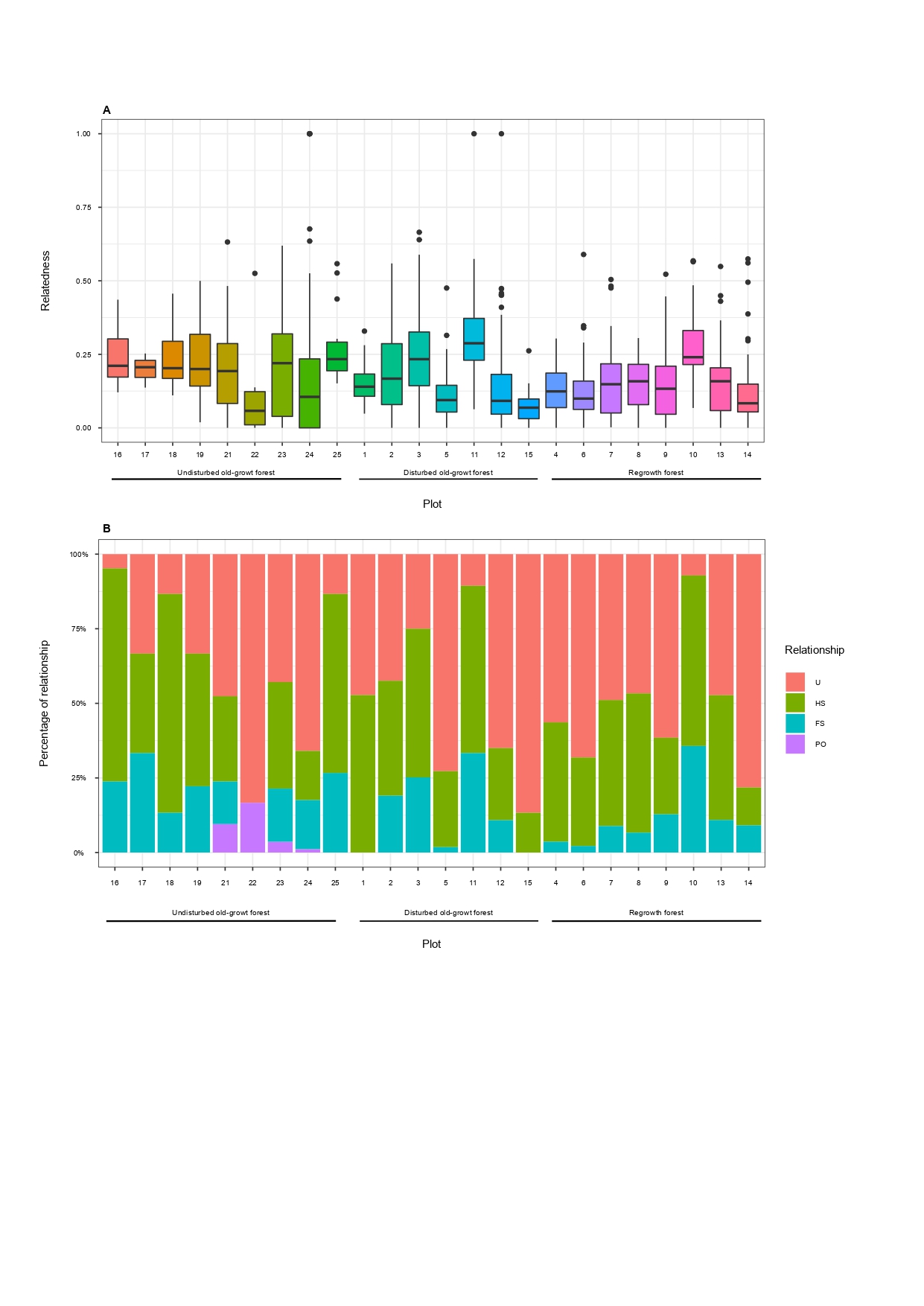
